# Deep learning permits imaging of multiple structures with the same fluorophores

**DOI:** 10.1101/2023.02.03.526797

**Authors:** Luhong Jin, Jingfang Liu, Heng Zhang, Yunqi Zhu, Haixu Yang, Jianhang Wang, Luhao Zhang, Yingke Xu, Cuifang Kuang, Xu Liu

## Abstract

Fluorescence microscopy is a powerful tool for life sciences, which employs fluorescent tags to label and observe cellular structures and their dynamics. However, due to the spectral overlap between different dyes, a limited number of structures can be separately labeled and imaged for live cell applications. Here we propose a novel double-structure network (DBSN) that consists of multiple connected models, which can extract six subcellular structures from three images with only two separate fluorescent labels. DBSN combines the intensity-balance models to compensate for uneven fluorescent labels for different structures and the structure-separation models to extract multiple different structures with the same fluorescent labels. Therefore, DBSN permits the imaging of multiple structures with only one fluorescent label. It significantly reduces photobleaching, breaks the bottleneck of the existing technologies, and would have vast applications in cell biology.

## 1 Introduction

Investigating the interactions between subcellular organelles has become one of the key research directions in cell biology. By using specific targeted fluorescent probes, researchers have been able to observe the distribution and dynamics of organelles in living cells with multi-color fluorescent microscopy imaging. Nevertheless, there are several limitations in monitoring of the organelle interactions by fluorescent microscopy, such as the low temporal resolution when using sequential imaging and the potential problems in spectral overlapping. In addition, it is impossible to ignore the switching time of filter or laser light source hardware devices in the study of multiple subcellular structures with sequential imaging.

With the development of microscopy imaging, especially super-resolution fluorescence microscopy technology, it is possible to discover more detailed observations of organelles. However, most of the existing super-resolution imaging technologies need large original data to achieve higher spatial resolution, which results in the cost of sacrificing time resolution. Structured illumination microscopy (SIM) [1], a representative super-resolution fluorescence microscopy technology that relies on delicate hardware devices to bypass the effect of point spread function (PSF), requires 9 or 15 raw images for 2-dimential (2D) or 3D imaging respectively. Computational algorithms have been designed to improve image resolution and achieve super-resolution reconstruction. For example, super-resolution radial fluctuations (SRRF) [2] is one of the most widely used computational super-resolution reconstruction methods, which requires hundreds to thousands of original data. The technologies with the highest spatial resolution, such as stochastic optical reconstruction microscopy (STORM) [3], photoactivated localization microscopy (PALM) [4], and DNA points accumulation for imaging in nanoscale topography (DNA-PAINT) [5], which accomplish super-resolution reconstruction based on single molecule localization algorithm requiring tens of thousands of raw data for a high spatial resolution image. Therefore, the time resolution of those super-resolution reconstruction methods is restricted by the excessive demand for raw data. In recent years, deep learning has been used in lots of studies to reduce the need for the large quantity of raw data [6-9], but achieves comparable spatial resolution. However, the damage of imaging delay becomes more serious when using the above super-resolution imaging technology to study organelle interaction under multi-channel sequence imaging.

Spectral overlap is another common problem in fluorescence microscopy. The typical method used in multi-structure imaging is to apply fluorescent probes with different excitation or emission spectra to carry out specific labeling of subcellular structures and then to combine with multi-channel imaging. Considering the overlapping between fluorescent probes, there are four different lasers commonly used in biological imaging research, 405 nm, 488 nm, 561 nm, and 640 nm. Thus, fluorescence imaging approaches are limited in the short number of different labels that can be distinguished in one image. Researchers have devoted to increasing the number of labels in a single cell. Lippincott-Schwartz and colleagues adopted a multi-spectral image acquisition method that overcomes the challenge of spectral overlap in the fluorescent protein palette. They have extended the number of different labels that can be distinguished in a single image into six [10]. Xi *et al* reported a spectrum and polarization optical tomography (SPOT) technique. Using Nile Red to stain the lipid membranes, SPOT can simultaneously resolve the membrane morphology, polarity, and phase from the three optical dimensions of intensity, spectrum, and polarization, respectively. These optical properties reveal lipid heterogeneities of ten different subcellular compartments [11]. To overcome the limited available labels in a single cell, both of them combined additional information such as spectrum or polarization in addition to image data. This will increase the complexity of the hardware system. Moreover, the SPOT technology is only suitable for objects with membrane structures and without specificity. Considering that fluorescent labeling may affect the function of cells, label-free imaging technology is also a hot topic of study. Several computational machine-learning approaches were proposed, which predicts some fluorescent labels from unlabeled biological samples [12-16]. However, these technologies require special imaging systems and are lacking universality. The complex background of bright field images will also reduce the accuracy of target structure extraction. In summary, these technologies mentioned above usually confronted with poor temporal resolution, fluorescence crosstalk, and limited by number of probes. Recently, deep learning-based algorithms have emerged that allow us to extract interesting information from a huge dataset without any prior knowledge.

Here, we propose a double-structure network (DBSN) via multiple connected networks, which can extract six subcellular structures from three images. The mechanism of the DBSN is mainly divided into three steps. Firstly, for each cell sample, three microscopic images are collected with three imaging channels. The bright filed image is used for segmentation of the nucleus and cell membrane. The fluorescence image of clathrin-coated pits (CCPs) and microtubules (MTs) is excited with a 488 nm laser and the fluorescence image of endoplasmic reticulum (ER) and adhesions is stimulated with a 561 nm laser. Then both the fluorescence images are pre-processed by intensity-balance models to compensate uneven distribution between two structures in a single image, which is typically caused by the diverse densities of fluorescent labels for different structures. Finally, the bright field image and the above two intensity-balanced images are fed into three different structure-separation models to extract the profile of the nucleus, cell membrane, CCPs, microtubules, ER, and adhesion (Fig. 1). In all, the proposed DBSN consists of five interconnected models that based on U-Net [17, 18]. The validated results demonstrate the value of the DBSN in improving the microscopy imaging speed and its significant applications for the imaging of dynamic interaction between organelles in cell.

**Fig. 1.**
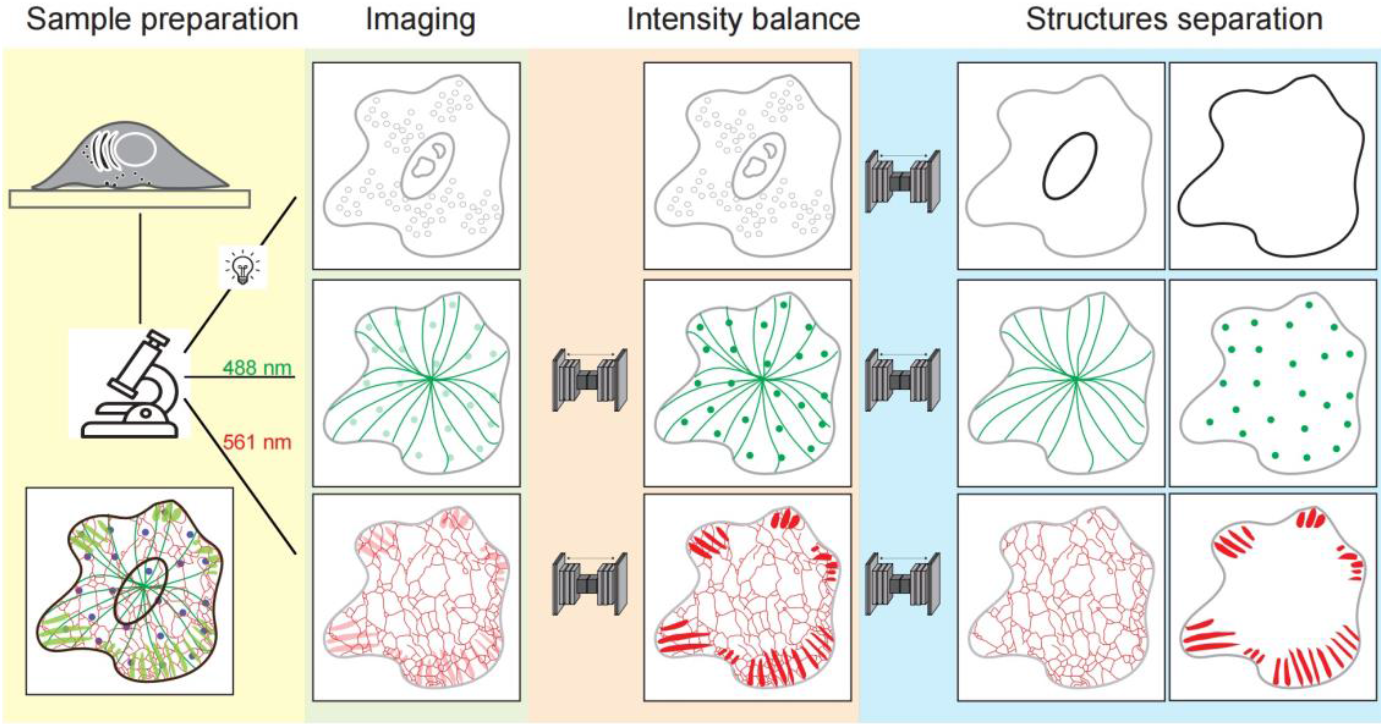
An overview of the double-structure network (DBSN). The DBSN consisting of three interconnected parts: imaging, intensity balance and structures separation. A bright field image is collected for nucleus and cell membrane segmentation. The clathrin-coated pits (CCPs) and microtubules (MTs) structures are labeled by clatherin-eGFP and EMTB-3×eGFP and image with 488 nm laser and the ER and cell adhesions are labeled with mCherry-KDEL and paxillin-mCherry and image with 561 nm laser. For each fluorescence channel, an intensity-balance model is adapted to compensate uneven intensity of the two structures in the same image. Finally, the bright field image of the same cell, together with the two intensity balanced images are fed into specific structure-separation models to extract six different subcellular structures.

## 2 The inspiration and pipeline to prepare training datasets for DBSN

Traditionally, to observe multiple different subcellular structures in the same cell at the same time, it is crucial to label these structures with different fluorophores and to image them in a single projective image. To alleviate the different z-axial distribution of different structures, it is important to compromise the focal plane to suppress the out-of-focus of certain structures (Fig. S1(a)). However, it is always challenging to ensure that the density of fluorophores in different structures are labeled at the same level. This directly leads to brightness discrepancies when displaying different subcellular structures in the same projective image (Fig. S1(b)). Typically, in imaging of multicolor structures, the resulting images with relatively balanced gray information among structures can be superimposed by adjusting the contrast in different channels separately. Such a method is inevitably a time-consuming procedure for multi-channel sampling. Therefore, we propose our method to segment multiple subcellular structures in the same cell with the same color labeling to achieve the purpose of synchronous imaging. In this case, the problem of intensity balance is particularly important in multi-structure images.

The fluorescence data of each subcellular structure was collected independently, whereas the training datasets used in this study were synthetic. Taking the two-dimensional spatial distribution of these four subcellular structures into consideration, we separated CCPs and MTs into one group, and ER and adhesion into another group. The pipeline to prepare training datasets can be divided into three steps (Figs. 2 (a) and (b)).

**Fig. 2.**
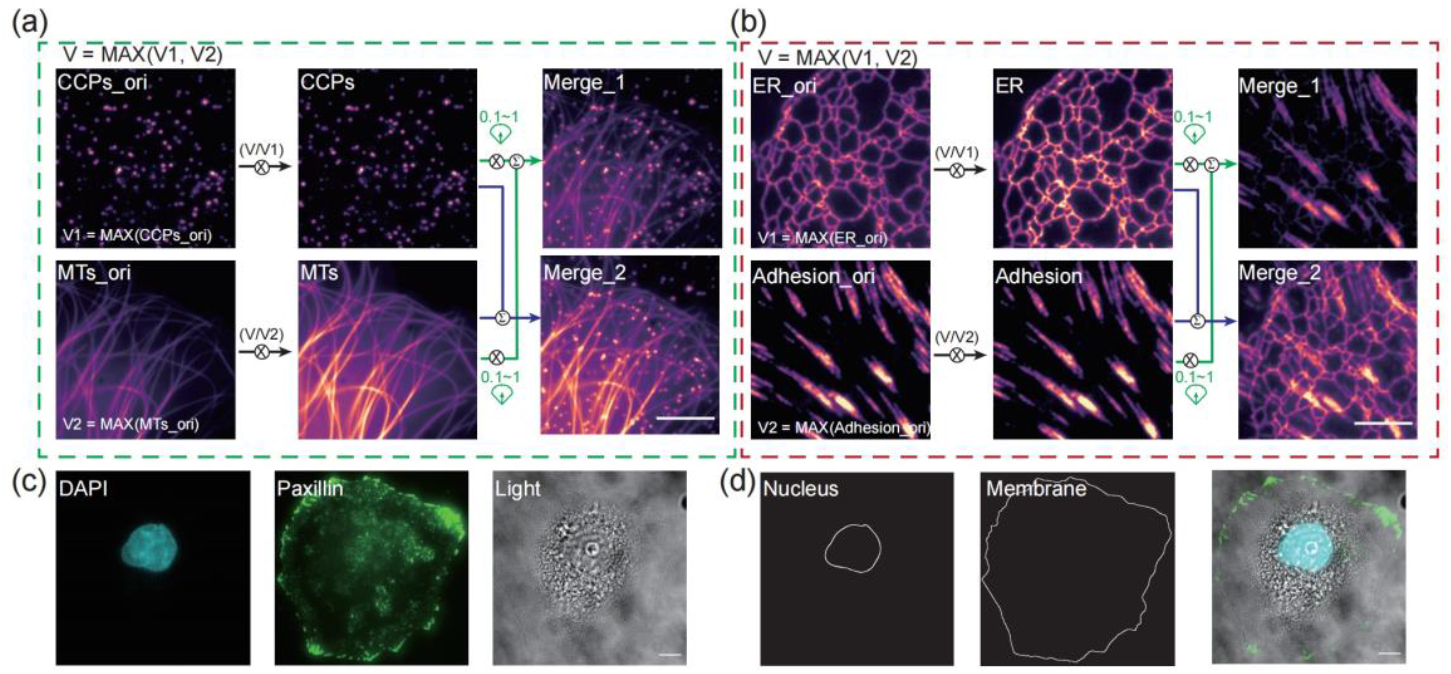
The pipeline to prepare fluorescence training datasets. Original fluorescence images named *CCPs_ori, MTs_ori, ER_ori* and *Adhesion_ori* are collected independently. (a) The maximum gray value *V* of *CCPs_ori* and *MTs_ori* are used to adjust the two images to the same intensity scale and to generate new images named *CCPs* and *MTs*. Two randomly generated weight coefficients between 0.1 and 1 are used to simulate an image with different intensities between the two structures (*Merge_1*), while the image named *Merge_2* is generated by directly adding the *CCPs* and *MTs* images. *Merge_1* is the input for the intensity-balance network. *Merge_2* is the ground truth for the intensity-balance network and the input of the structure-separation network. *CCPs* and *MTs* are the ground truth for the structure-separation network. (b) The pipeline to prepare ER and adhesion datasets is similar as described in (a). (c) A Hoechst 33342 labeled nucleus image, a paxillin-eGFP labeled cell adhesion image, and a bright field view of the cell are collected respectively. The bright field image is used as the input for the structure-separation network. (d) The profiles of the nucleus and cell membrane are manually segmented, which are then used as the ground truth of the structure-separation network. Scale bars: (a), (b) 5 μm, (c), (d) 10 μm.

For each fluorescence sample, consider two 2D images { *I*_*1*_, *I*_*2*_} ∈ ℝ^*m×n*^contained enough structures information. First, we searched the maximum gray value of *I*_*1*_ and *I*_*2*_. Then, this maximum gray value was used to pull the two images to the same intensity scale named *I*_*1_gt*_and *I*_*2_gt*_. That is illustrated as:

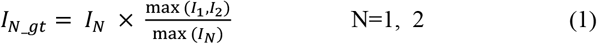

Then, randomly generated two weight coefficients between 0.1 and 1, and multiplied them with the above two intensity balanced images to prepare images with random intensity difference:

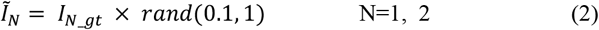

An intensity-balance network attempts to compensate the diverse densities of fluorescent labels for different structures by finding a function *f*_θ :_ℝ^*m×n*^→ℝ^*m×n*^(parameterized by the network weights *θ*) such that

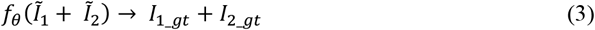

Finally, a structure-separation network attempts to extract two different subcellular structures from the intensity-balanced summed image by finding a function *g*_δ_ : ℝ^*m×n*^→ℝ^*m×n*^(parameterized by the network weights δ) such that

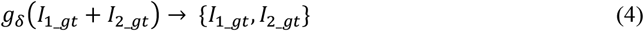

To further reduce the complexity of fluorescent labeling, the bright field image was used to segment the nucleus and cell membrane (Fig.2 (c)). For the same cell, Hoechst 33342 labeled nucleus, Paxillin-GFP labeled adhesion, and bright field images were captured separately. We used the bright field data as the input for the structure-separation network, whereas the Hoechst 33342 and Paxillin labeled images were manually segmented as the ground truth (Fig. 2 (d)). Detailed information about each dataset is listed in Table S1.

## 3 Intensity-balance network for multi-structure imaging

The architecture of the intensity-balance network is described in Fig. 3(a). The fluorescent dual-structure image with intensity difference is the input of the network. The intensity-balanced image is the ground truth of the network (see equation (3)). The local enlargement views shown represent the structure information vanished caused by the intensity difference, which can be restored by the intensity-balance network (Fig. 3(a)).

**Fig. 3.**
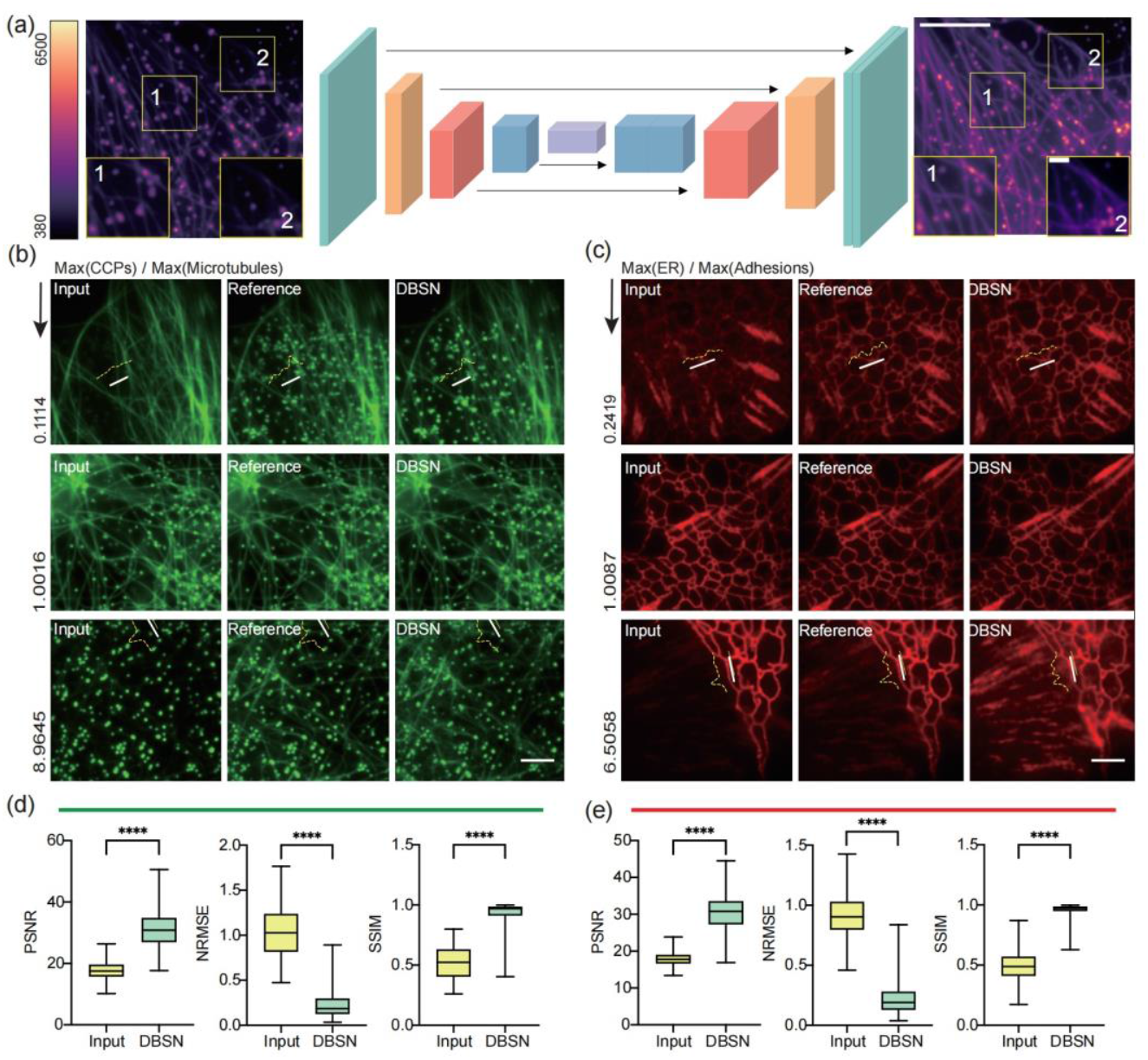
The architecture of intensity-balance network and its performance evaluation. (a) The architecture of the intensity-balance network. The fluorescent dual-structure image with intensity difference is the input of the network. The intensity-balanced image is the ground truth of the network. (b, (c) Side-by-side comparison of the intensity-balanced results between the ground truth and DBSN under different situations. For each panel, the first column shows the input data with different intensity ratios of the two structures. The intensity difference ratio is marked at the bottom left of each input image. The second column shows the ground truth, whereas the last column is the output of the network. The yellow dashed lines illustrate the single pixel intensity distribution of the white lines in each corresponding images. (d), (e) Quantitative results of intensity-balance models as evaluated by PSNR, NRMSE and SSIM respectively. Scale bars: (a) 5 μm, 1 μm, (b), (c) 3μm. ^****^ represents *p*<0.0001.

The recovered results by using the intensity-balanced network were shown in Figs. 3(b) and (c). For each panel, the first column is the input data with different intensity ratios of the two structures. Notice that when the maximum intensity value of the CCPs image is only one-tenth of the maximum intensity value of the microtubule image, intuitively, we can hardly see the CCPs in the input image (Fig. 3(b)). A similar situation can also be observed in the combination diagram of ER and adhesion. When the maximum intensity value of ER image is only one-fifth of the maximum intensity value of the adhesions image, we can barely distinguish the shape of ER from the input image (Fig. 3(c)). Similarly, when the intensity of CCPs or ER is several times higher than the other structures, the information of them almost disappears in the input image. All the reference images shown represent the manually intensity-balanced dual structure images, while the last columns in the Figs. 3(b) and (c) are the output of the network. The yellow dashed lines on the images are intensity distribution of the single pixel white lines in each image. Compared with the input image, the shape of the intensity distribution curve in the DBSN image is closer to the ground truth images and contains more information.

Finally, we statistically evaluated the performance of intensity-balance models with parameters peak signal-to-noise ratio (PSNR), normalized root-mean-square error (NRMSE), and structural similarity (SSIM) by using manually adjusted intensity-balanced images as references (Figs. 3(d) and (e)). The PSNR, NRMSE, and SSIM performance confirms that DBSN can significantly balance the intensity information in the input images. Those three parameters were also used to evaluate the performance of the intensity-balance network training procedure (Fig. S2). We measured them for every 10 epochs, and based on the performance we selected the optimal intensity-balance model.

## 4 Structure-separation network for multi-structure segmentation in a single image

After intensity normalization, multiple subcellular structure images were superimposed into a projection image as the input of the structure-separation network, and the single structure microscopic image was used as the ground truth. The value of PSNR, NRMSE, and SSIM were calculated to evaluate the performance of the structure-separation network training procedure (Fig. S3). We measured those three parameters for every 10 epochs, and selected the optimal structure-separation model based on their performance.

The efficacy of the structure-separation network is presented in Fig. 4. As shown in Figs. 4(a)-(d), from left to right are the input images, the ground truth of these two structures, and network separation results, respectively. The input image is a dual structure overlayed image after manually intensity balanced. As shown in Fig. 4(b), both CCPs and MTs are observed in the input image. The two single-structure ground truth images showed that there are bright CCPs structure superimposed on the end of the microtubule indicated by the yellow dotted line. DBSN distinguishes and segments these two structures perfectly. In the output results from the DBSN, only CCPs are retained in the channel of CCPs, and only the microtubule structure in the channel diagram corresponding to microtubule structure. The fluctuation of the gray value of the corresponding positions in the two images is very similar to the reference images. Similarly, as shown in the local enlargement images in Fig. 4(d), both ER and adhesions are merged in the input image, which is difficult to distinguish these two structures visually. Compared with the ground truth images, the output of DBSN provides comparative reliability, which can be further validated by the quantitative evaluation results shown in Fig. 4(e).

**Fig. 4.**
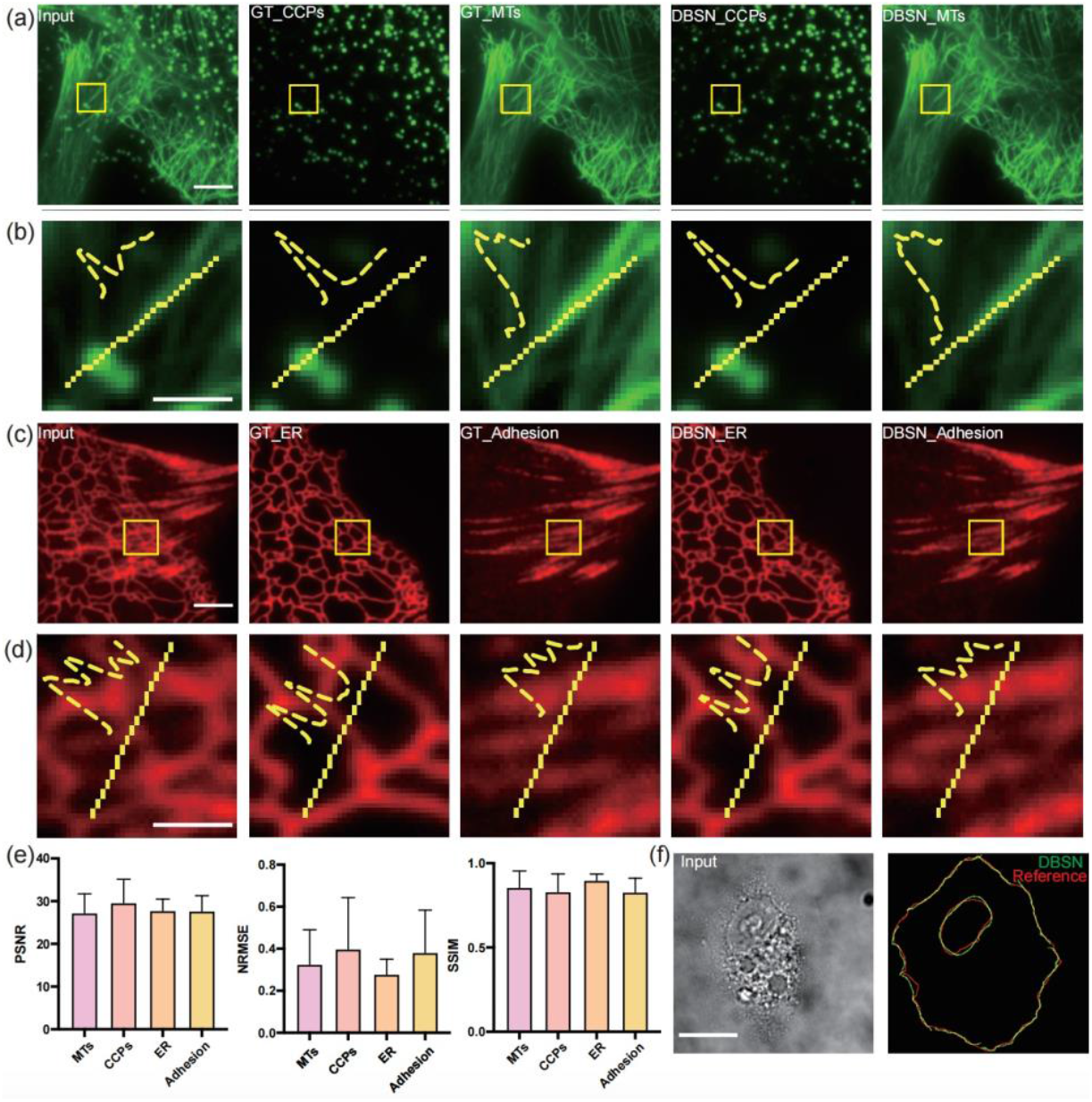
The performance of DBSN in multiple structure separation. (a) Separation of the CCPs and MTs merged image. From left to right are input image, ground truth (GT) of the two structures and DBSN separated results. (b) Enlarged regions enclosed by the yellow boxes in (a) with intensity distribution profiles along the yellow dashed lines. (c) Separation of ER and adhesions overlayed image. From left to right are input image, GT of the two structures and DBSN results. (d) Enlarged regions enclosed by the yellow boxes in (c) with intensity profiles along the yellow dashed lines. (e) Quantitative evaluation results of structure-separation models by PSNR, NRMSE and SSIM analyses. (f) Nucleus and cell membrane segmentation results. Left is the input bright field image. Right shows the segmentation results. The red curves show the manually marked reference and the green curves are the output of the network. Scale bars: (a), (c) 3μm, (b), (d) 1 μm, (f) 10 μm.

In Fig. 4(f), the left image is the input bright field image, and the right one marks the manually labeled reference results of the cell nucleus and plasma membrane (red circles) and the corresponding output of the DBSN (green circles). Our results demonstrated that DBSN identifies these two structures perfectly as the network output results matched correctly in spatially with the ground truth.

## 5 Structure separation ability of DBSN on live cell imaging application

We next investigated the capability of DBSN for use in live cell imaging. Since deep learning models are sensitive to the data distributed characters, it is not wise to apply the previous models to the images from a different device. Transfer learning provides deep learning with better transferability, economizes the training datasets and allows researchers to deal with different but similar problems easily. Therefore, using the NCF950 microscope provided by Novel Optics, we recorded and organized a new training dataset to train more reliable models (Table S2).

Firstly, the pre-trained intensity-balanced models were used to initialize two new U-Net based intensity-balance models. As demonstrated in Fig. S4, by using the new dataset ({*Î*_*1*,_ *Î*_*2*_} ∈ ℝ^*m×n*^) to retrain the models, we acquired the considerable efficient intensity-balance model 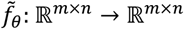 (Fig. S4). After that, all of the training data were refined by those intensity-balance model to prepare the dataset for the next structure-separation models 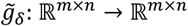. Resemble the first step, the pre-trained structure-balanced models were used to initialize two new U-Net based structure-balanced models (Fig. S5).

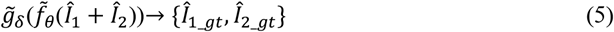

For live cell applications, the COS-7 cells were co-transfected with clathrin-eGFP, EMTB-3×eGFP, mCherry-KDEL and Paxillin-mCherry plasmids to label the CCPs, MTs, ER and adhesions respectively. Time-lapse fluorescence images were collected with the NCF950 microscope. As shown in Fig. 5, when two subcellular structures in the same cell were simultaneously imaged with the same fluorophores, DBSN can effectively separate different structures into different channels (Video 1, MP4, 5.6 MB; Video 2, MP4, 905 KB; Video 3, MP4, 5 MB;).

**Fig. 5.**
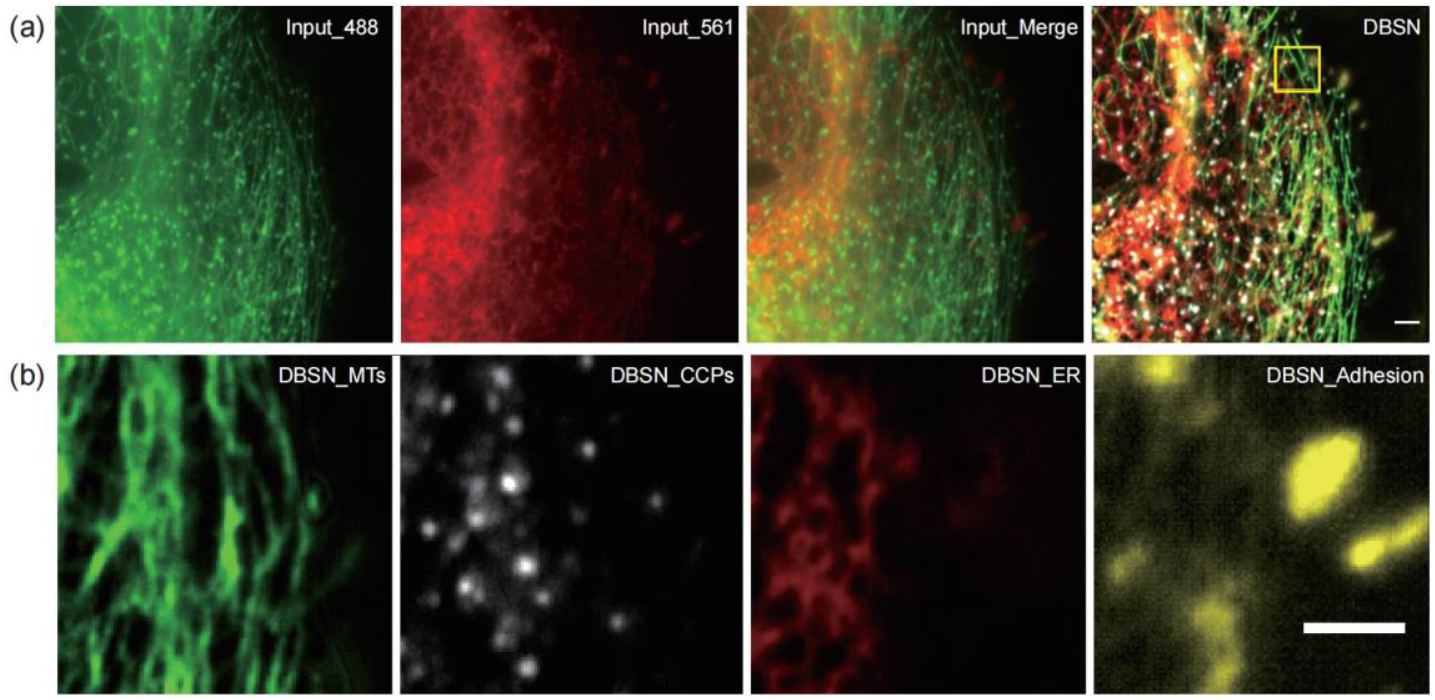
The DBSN separates different structures in a cell with the same fluorophore labeling. The COS-7 cell is expressing clatherin-eGFP, EMTB-3×eGFP, mCherry-KDEL and paxillin-mCherry. Only a single time point is shown for two channels and the DBSN processed result. (b) Higher magnification of the rectangle area in (a), which highlights the reliability of DBSN in details (Video 2, MP4, 905 KB). Scale bars: 5 μm.

## 6 Discussion and Conclusion

This study focuses on the realization of rapid imaging of the dynamic processes of the interactions between multiple subcellular structures in living cells. Through deep learning based neural network to separate multiple structures with the same labeling, our proposed method will increase acquisition speed and solve the problem of limited types of fluorophores available for multi-color imaging. By using the proposed DBSN method, one could extract six subcellular structures from only three microscopic images of the same cell. The established intensity-balance model addresses the challenge that different labeling density or expression levels of the fluorescent markers between different structures may lead to significant intensity differences in the acquired projective image. Whereas the structure-separation models overcome the restriction of only four different structures can be separately imaged with fluorophores available with distinct spectra in our microscope setup. Based on the DBSN, multiple subcellular structures can be labeled with the same color and then imaged simultaneously. It would truly achieve the real-time recording of the interactions between different subcellular organelles and facilitate the studying of their regulatory mechanism. According to the current study, with new training datasets and by optimization of the deep learning network, we anticipate that it would be feasible to separate more than two structures labeled with the same fluorophore at the same time.

Moreover, finding the solution to handle the problem of defocusing between subcellular structures (Fig. S1) will further improve the performance of DBSN. A cycle generative adversarial network-based model and a multi-component weighted loss function have been used to solve the out-of-focus issue in microscopy [19]. If our DBSN method integrates the defocus correction model, it will promote the reliability of multi-structure imaging. In addition, by taking the microscopic images acquired on other imaging modalities, such as spinning-disc confocal SIM (SD-SIM) [20], STED [21], two-photon microscopy [22], 3D SIM [23], and expansion microscopy [24] into the training datasets, the application of our developed DBSN method could be further expanded. Some of the training images of this study were from the public platform BioSR [25]. However, there is still a lack of open-source public datasets for microscopy image analysis. Even a slight difference in imaging procedure will degrade the performance of trained models. Using synthetic datasets or public datasets in the pretraining step and then combining transfer learning on the existing data is an optimized solution to reach satisfactory results. As deep learning technologies developed, an intelligent augmented microscope [26] and event-driven microscopes [27, 28] have already attracted the interests of biomedical researchers. However, all those work rely on a huge quantity of annotated datasets with high quality. Thus, it is important to establish a public platform for microscopy images with credibility.

## Acknowledgments

This work was supported by the National Natural Science Foundation of China (62105288 and 22104129), Zhejiang Provincial Natural Science Foundation (LQ22F050018 andLZ23H180002), National Key Research and Development Program of China (2021YFF0700305), Zhejiang University K.P.Chao’s High Technology Development Foundation (2022RC009), the Fundamental Research Funds for the Central Universities (226-2023-00091) and Fellowship of China Postdoctoral Science Foundation (2021M692831).

## Code and Data Availability

All the training datasets are available from the corresponding author upon request, due to size limitations. The source codes for PyTorch and part of the source image datasets have been uploaded to GitHub: https://github.com/luhongjinzju/DBSN. Other relevant data are also available from the corresponding author upon request, due to size limits.

**Luhong Jin** received her PhD from Zhejiang University at 2020. Currently, she is a postdoc at the Department of Biomedical Engineering at Zhejiang University. She was a visiting PhD student at the Department of Pharmacology, University of North Carolina at Chapel Hill, from 2018 to 2019. Her research interests are fluorescent super-resolution microscopy imaging and its information analysis.

**Yingke Xu** received his BS and PhD degrees from Zhejiang University at 2003 and 2008, respectively. From 2008 to 2012, he worked initially as a Postdoctoral Fellow at Yale University School of Medicine, and was subsequently promoted to be an Associate

Research Scientist at the Cell Biology Department at Yale University. Start from 2014, he is a Visiting Associate Professor at Yale University and an honorary visiting scientist at the laboratory of Dr. James Rothman (2013 Nobel Laureate in Physiology or Medicine) at ShanghaiTech University. Now, he is a professor at the Department of Biomedical Engineering at Zhejiang University, with research interests in optical super-resolution imaging and cell biology studies.

Biographies of the other authors are not available.

