## Supplementary information for "Deep learning permits imaging of multiple structures with the same fluorophores"

### 1. Cell culture and datasets preparation

The COS-7 cells were cultured in high glucose Dulbecco's modified Eagle's medium (DMEM) (Gibco, #11965092), supplemented with 10% fetal bovine serum (Sigma-Aldrich, #F8313) and 1% penicillin–streptomycin (Beyotime, #C0222) at 37°C in a humidified 5% $CO_2$ incubator. Cells were planted into a 35-mm glass bottom dish (Cellvis, #D35-20-1-N) for fluorescence imaging experiments. For live cell imaging study, the COS-7 cells were transfected with clatherin-eGFP, EMTB-3×eGFP, paxillin-mCherry, and mCherry-KDEL to label the CCPs, MTs, cell adhesions and ERs respectively.

For the training and testing step, original SIM raw images of CCPs, MTs, and ER were downloaded from the BioSR dataset [1]. Original adhesion SIM raw images were from our previous work [2]. The average images of all the above SIM raw images were used in training of the DBSN. To visualize and segment the plasma membrane, COS-7 cells were transfected with paxillin-eGFP. The cell nucleus was stained with Hoechst 33342 (Thermo Scientific, #62249). For each cell, a nucleus image, an adhesion image, and a bright field image were collected. Those three images were used to train the nucleus and cell membrane segmentation model. For the transfer learning step, the COS-7 cells co-transfected with clatherin-mCherry and EMTB-3×eGFP were used, and the acquire images were for training the intensity-balance and structure-separation models for CCPs and MTs. The COS-7 cells co-transfected with mCherry-KDEL and paxillin-eGFP were used to acquire samples to train the intensity-balance and structure-separation models for ER and adhesions.

### 2. Microscopes

NCF950 fluorescence microscope made by Novel Optics (Ningbo, China) is equipped with 405 nm, 488 nm, and 561 nm laser lines. The objective lens is 100X (1.45 NA) and the CMOSE camera is HAMAMATSU C13440. For Hoechst 33342-labeled nucleus imaging, a 405-nm laser (5mW) was used as excitation light and for visualization of paxillin-mCherry labeled adhesions, a 561-nm laser (100 mW) was used for imaging. The widefield images were acquired under the control of Novel Optics software with an exposure time of 40 ms.

### 3. Image pre-processing

For training the intensity-balance models and fluorescence structure-separation models, we cropped the original image stacks into small patches to generate more training samples. Images of CCPs, MTs, adhesions, and ER were cropped into 256 × 256 pixels patches. To prepare the datasets for the nucleus and cell membrane segmentation model, the whole cell should be contained in the image. Therefore, instead of cropping patches, we resized the original image into 256 × 256 pixels. After manually segmented the profiles of the nucleus and cell membrane, image flipping, rotating, and mirroring were used to augment the datasets. In total, we obtained 600~800 samples for each experiment, which were then randomly divided into training and testing subsets. Detailed information about each dataset is listed in Table S1.

For transfer learning, we cropped the original image stacks into 512 × 512 pixels. Image flipping, rotating, and mirroring were used to augment the datasets. In total, we obtained 600~700 samples for each experiment. Among them, 50 samples were used as testing subsets. Detailed information about each dataset is shown in Table S2.

### 4. Training procedures

We adopted the Python package U-Net as previously reported [3]. The training and inference were performed on a computation platform with Intel Core i9-10900KF CPU and graphic processing cards (NVIDIA) GeForce RTX 3080 GPU. During the training process, we used the Adam optimizer. We initialized the networks randomly and trained the models with a typical starting learning rate of $1 \times 10^{-4}$. We trained the intensity-balance network and structure-separation network for 5,000 epochs, and saved the models for every 10 epochs. The representative plots for validation of the PSNR, NRMSE, and SSIM during the training processes of different networks are shown in Fig. S2 and Fig. S3. The codes for training and testing were written using Python with PyTorch framework. The PSNR, NRMSE, and SSIM measurements were performed as previous described [2]. All the source codes are free available online (https://github.com/luhongjinzju/DBSN).

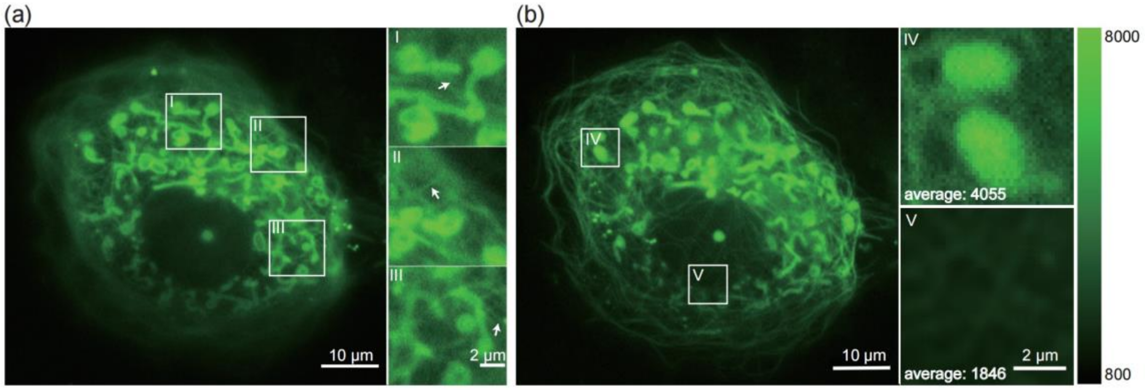


**Fig. S1.** Representative images show the defocusing and uneven expression of fluorescent proteins for dual-structure microscopic imaging with the same fluorophore. The COS-7 cells were transiently transfected with Tom20-eGFP and EMTB-3×eGFP. (a) Due to the different z-axial distribution of different structures, some objects in the two-dimensional projection image are out of focus. Enlarged views in I-III represent several examples of mitochondria that are in the right focus but MTs structure with different degrees of defocusing. (b) Owing to transient transfection, the expression levels of the fluorescent markers in different structures are different, which leads to a large difference in intensity in acquired image. Enlarged images in IV and V demonstrate two local areas with significant intensity differences.

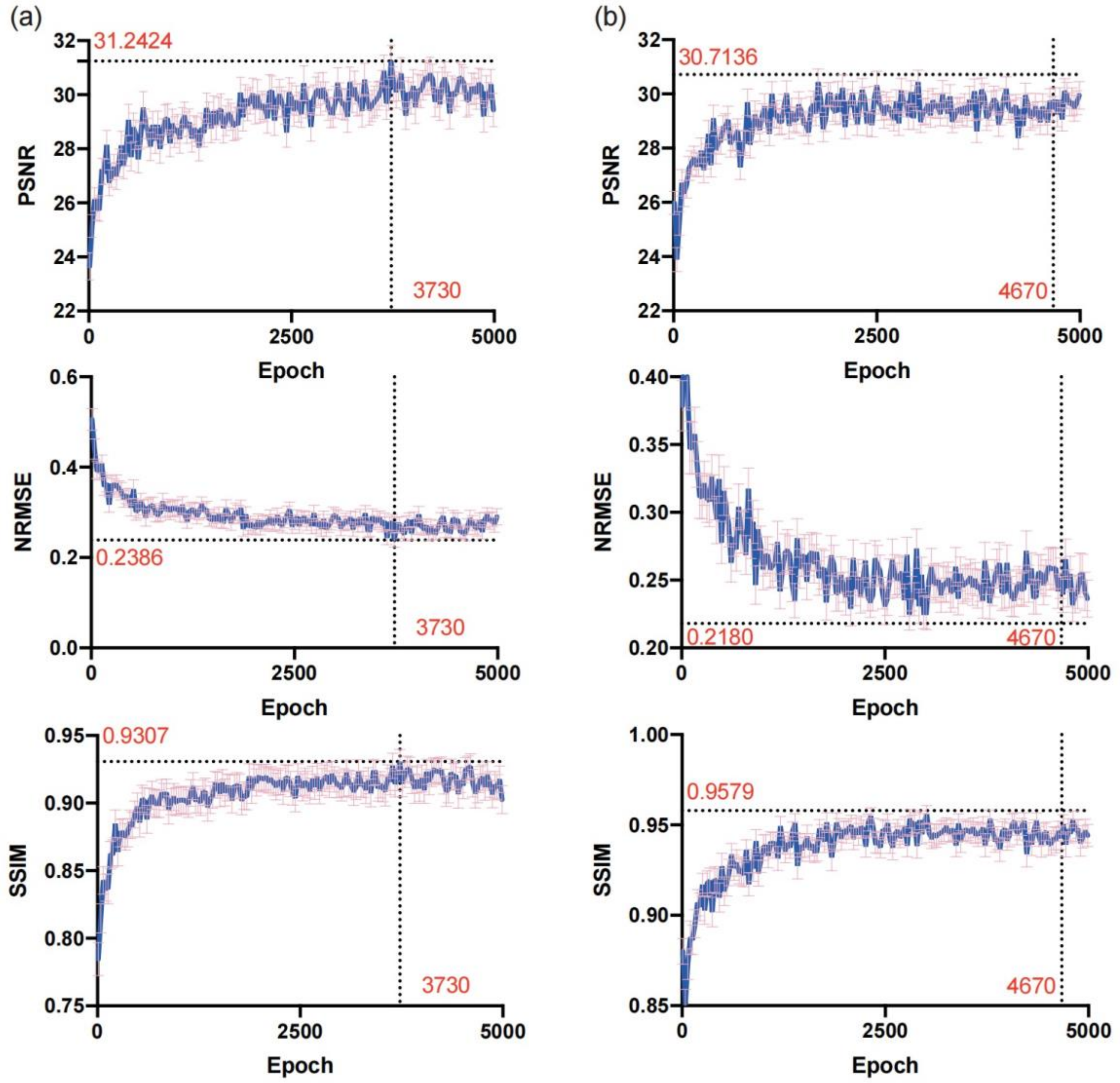


**Fig. S2. The performance of the intensity-balance network monitored during the training process.** (a) The distribution of PSNR, NRMSE, and SSIM analyses during the training of CCPs and MTs intensity-balance model. The model with epoch=3730 is selected. (b) The distribution of PSNR, NRMSE, and SSIM during the training of intensity-balance model for ER and adhesions. The model with epoch=4670 is selected.

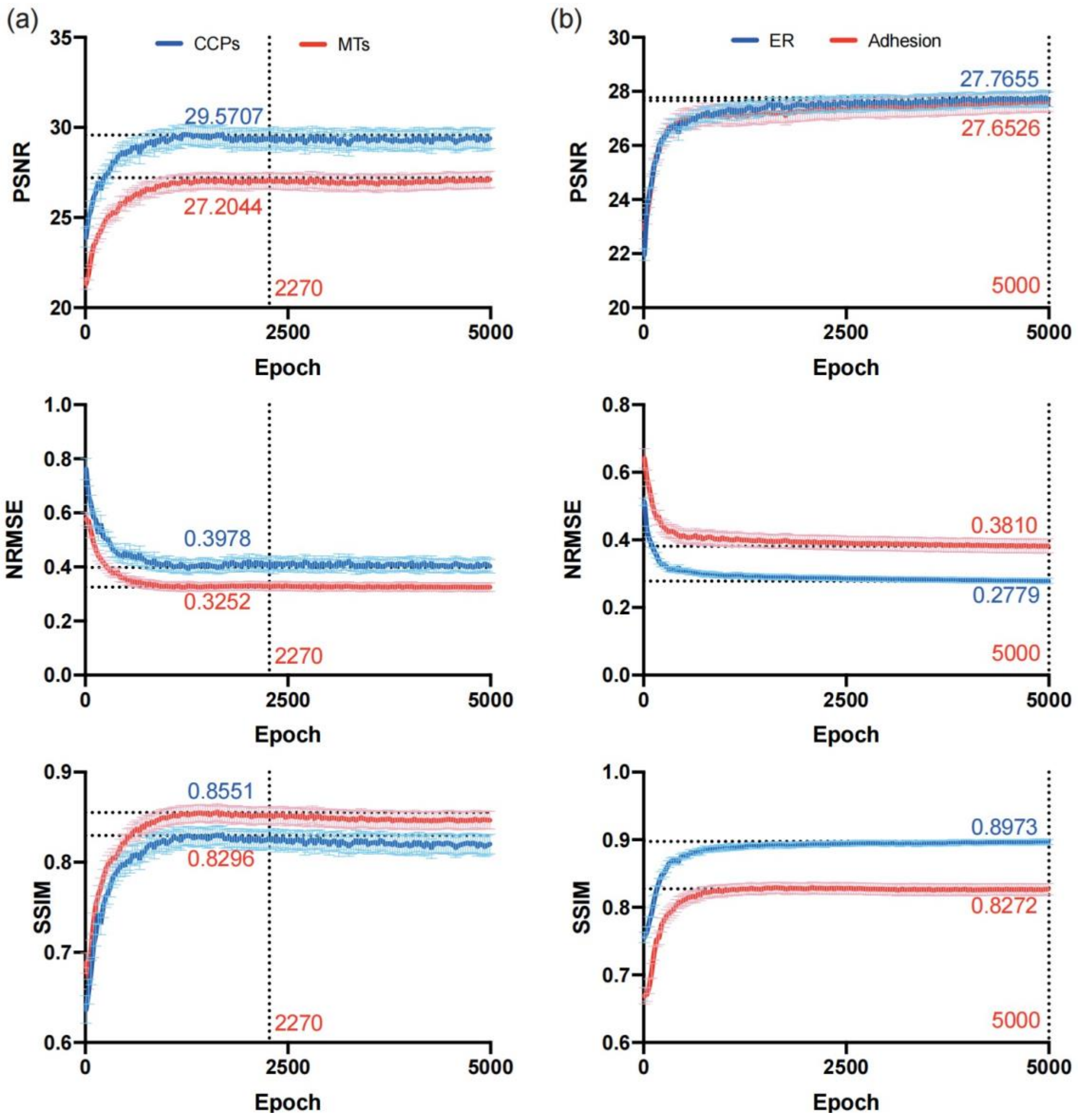


**Fig. S3. The performance of structure-separation network analyzed during the training process.** (a) The analyses of PSNR, NRMSE, and SSIM during the training of CCPs and MTs structure-separation model. The model with epoch=2270 is selected. (b) The distribution of PSNR, NRMSE, and SSIM during the training of structure-separation model for ER and adhesions. The model with epoch=5000 is selected. At this point, there is no significant improvement in model performance.

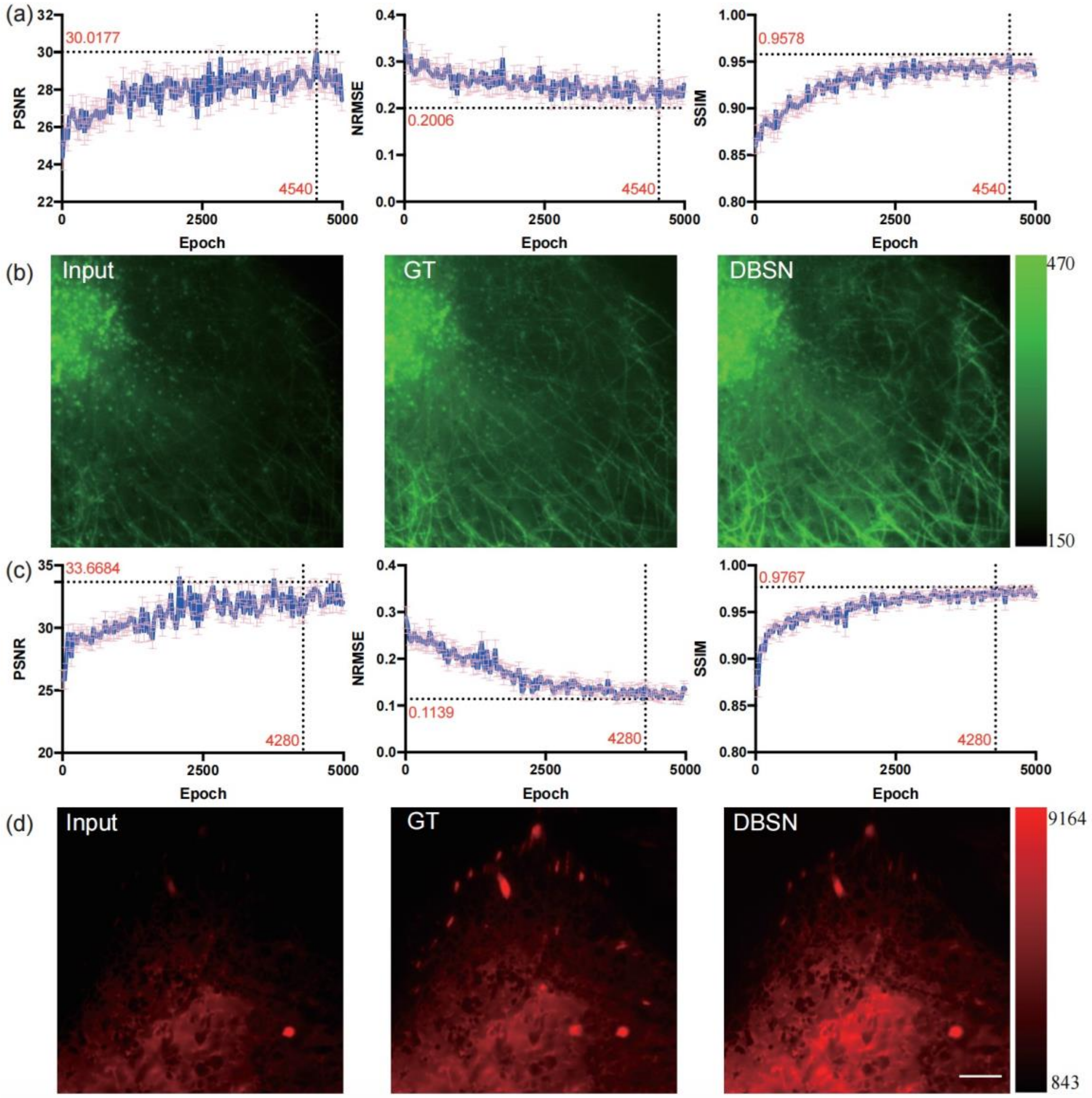


**Fig. S4. Validation the transfer learning performance of the intensity-balance network.** (a) and (c) The performance of transfer intensity-balance network is illustrated by PSNR, NRMSE, and SSIM distributions during the training process. For CCPs and MTs, the model with epoch=4540 is selected. For ER and adhesions, the model with epoch=4280 is selected. (b) and (d) Representative intensity-balanced results. From left to right are input image, ground truth (GT) and the output of the network result. Scale bars: (b), (d) 5 μm.

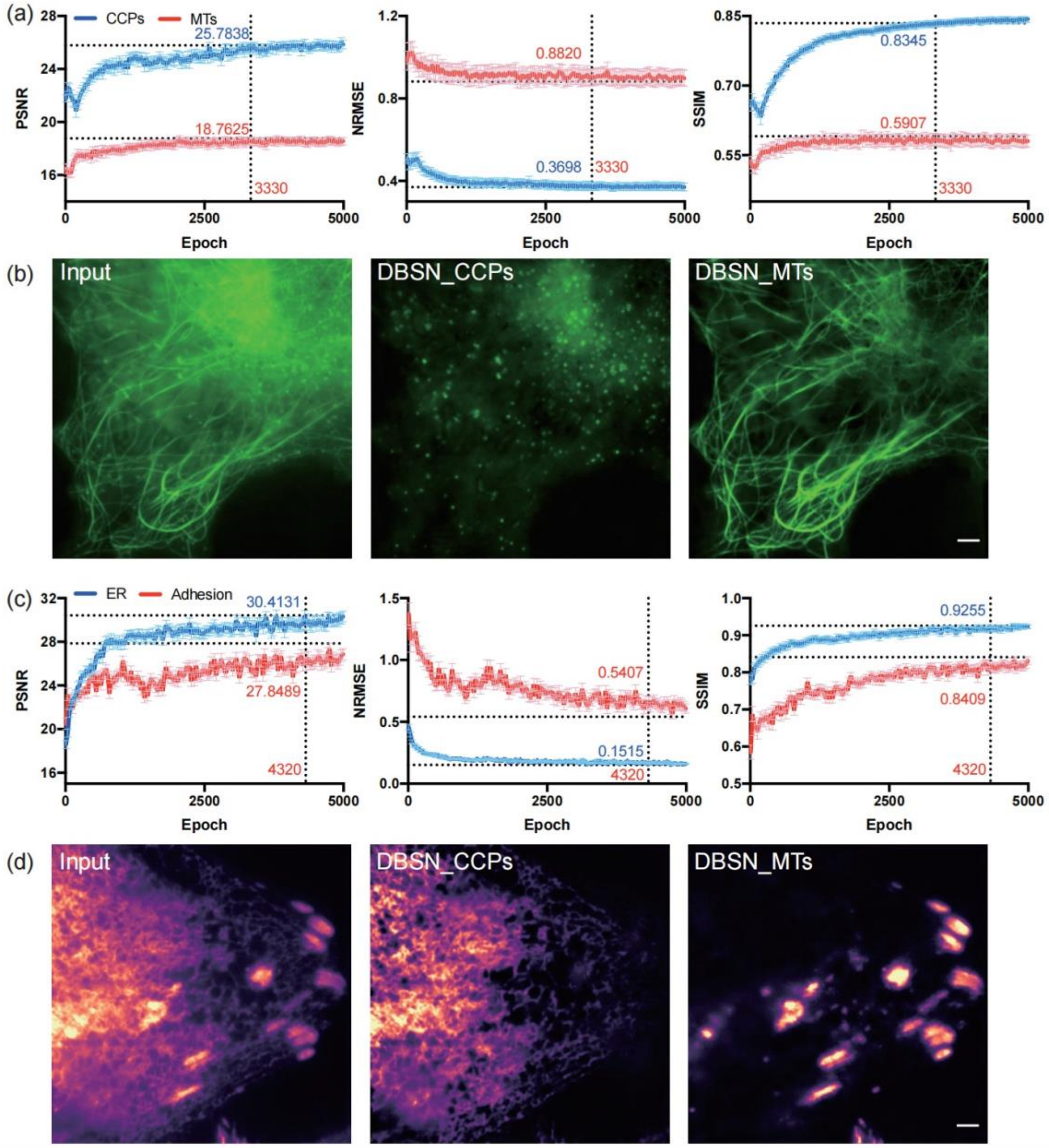


**Fig. S5. Validation the transfer learning performance of the structure-separation network.** (a) and (c) The performance of transfer structure-separation network is analyzed by PSNR, NRMSE, and SSIM distributions during the training process. For CCPs and MTs, the model with epoch=3330 is selected. For ER and adhesions, the model with epoch=4320 is selected. (b) and (d) Representative structure-separation results. Scale bar: (b), (d) 5 μm.

**Table S1. The size of datasets for training and testing in each model.**

| | Original images | Training patches (256 × 256) | Testing patches (256 × 256) |
|---|---|---|---|
| Nucleus and cell membrane | 109 | 719 | 16 |
| CCPs and MTs | 200 | 720 | 109 |
| ER and adhesions | 200 | 558 | 96 |

**Table S2. The size of datasets for training and testing in each transfer model.**

| | Original images | Training patches (512 × 512) | Testing patches (512 × 512) |
|---|---|---|---|
| CCPs and MTs | 170 | 676 | 50 |
| ER and adhesions | 145 | 555 | 50 |

**Video 1** Description: The DBSN separates different structures in a living cell. Shown are the widefield merged structures images and the output of DBSN. Scale bar: 5 μm.

**Video 2** Description: The detailed local cropped movies of DBSN prediction. Shown are the prediction of MTs, CCPs, ER and adhesions. Scale bar: 5 μm.

**Video 3** Description: The DBSN in time-lapse imaging of MTs, CCPs, ER and adhesions labeled with two identical fluorophores.

*References*


1. C. Qiao et al., "Evaluation and development of deep neural networks for image super-resolution in optical microscopy," *Nat. Methods* **18**(1), 194-202 (2021).
2. L. Jin et al., "Deep learning enables structured illumination microscopy with low light levels and enhanced speed," *Nat. Commun.* **11**(1), 1934 (2020).
3. O. Ronneberger, P. Fischer, and T. Brox, "U-Net: convolutional networks for biomedical image segmentation," *International Conference on Medical Image Computing and Computer Assisted Intervention (MICCAI)* 234–241 (2015).
